# Toll-like receptor signalling via IRAK4 confers epithelial integrity and tightness through regulation of junctional tension

**DOI:** 10.1101/2021.10.25.465823

**Authors:** Jesse Peterson, Kinga Balogh Sivars, Ambra Bianco, Katja Röper

**Affiliations:** MRC-Laboratory of Molecular Biology, Francis Crick Avenue, Cambridge Biomedical Campus, Cambridge CB2 0QH, UK; Clinical Testing and Precision Medicine, Global Procurement, Operations & IT, AstraZeneca, Gothenburg, Sweden; Clinical Pharmacology and Safety Sciences CPSS | Oncology Safety, AstraZeneca, Darwin Building, Cambridge Science Park, Milton Road, Cambridge CB4 0WG, UK

**Keywords:** Toll-like receptors, intestinal epithelium, epithelial barrier, tight junctions, cytoskeleton, tension

## Abstract

Toll-like receptors (TLRs) in mammalian systems are well characterised for their role in innate immunity. In addition, TLRs also fulfil crucial functions outside immunity, including the dorso-ventral patterning function of the original Toll receptor in *Drosophila* and neurogenesis in mice. Recent discoveries in flies suggested key roles for TLRs in epithelial cells in patterning of cytoskeletal activity near epithelial junctions. Here we address the function of TLRs and the downstream key signal transduction component IRAK4 (interleukin-1 receptor <u>a</u>ssociated <u>k</u>inase <u>4</u>) in human epithelial cells. Using differentiated human Caco-2 cells as a model for the intestinal epithelium, we show that these cells exhibit baseline TLR signalling as revealed by p-IRAK4 and that blocking IRAK4 function leads to a loss of epithelial tightness involving key changes at tight junctions and adherens junctions. These changes correlate with a loss of epithelial tension and changes in junctional actomyosin. Knock-down of IRAK4 and certain TLRs phenocopies the inhibitor treatment. These data suggest a model whereby TLR receptors near epithelial junctions might be involved in a continuous sensing of the epithelial state to promote epithelial tightness and integrity.

## Introduction

Epithelia are one of the major tissue types in all animals and provide the building blocks for most organs in both invertebrates and vertebrates. Furthermore, organ morphogenesis during development in many cases commences from initially simple epithelial primordia. Epithelia also represent the first barrier of defence against infection and help mount an innate immune response.

Toll-like receptors (TLRs) are key components of the innate immune response, with their activation through pathogen-related molecules leading to transcription and production of e.g. cytokines (Leulier and Lemaitre, 2008; Netea et al., 2012). Most TLRs are single pass transmembrane receptors with a leucine-rich extracellular region (LRR) that recognizes pathogen-associated molecular patterns (PAMPs) on bacteria, viruses, fungi and other unicellular pathogens. This pathway is conserved from invertebrates to mammals. The activated receptor recruits to its intracellualr tail the adaptor MyD88 which through its death domain binds to the downstream kinases IRAK4 (interleukin-1 receptor <u>a</u>ssociated <u>k</u>inase <u>4</u>) and then IRAK1 or 2 (interleukin-1 receptor <u>a</u>ssociated <u>k</u>inase <u>1/2</u>) to form the so-called myddosome. Through further downstream components this eventually leads to degradation of cytoplasmic IκB (inhibitor of nuclear factor κ-B), releasing NFκB (nuclear factor κ-light-chain-enhancer of activated B cells) to activate targets in the nucleus (Netea et al., 2012). NFκB-dependent gene transcription regulates a variety of targets including factors that are directly involved in the immune response such as cytokines and chemokines, but also cell adhesion molecules, growth factors and their receptors and apoptosis related factors (http://www.bu.edu/nf-kb/gene-resources/target-genes/). TLR signalling has been implicated in a significant number of immune disorders, and therefore members of this pathway, such as TLRs, IRAK kinase family members, IKKs, have come to attention as promising drug targets.

Toll-like receptors were first identified in *Drosophila* where, similar to mammals, several TLRs exist. In the fly, where their role has been extensively investigated, important functions during innate immune responses but also during development have emerged. In fact, the receptor Toll itself was initially characterised during the process of dorso-ventral axis formation in the fly embryo, where localised receptor activation leads to the nuclear localisation of one of the fly NFκBs, Dorsal, in only the future dorsally located cells (Gerttula et al., 1988). Further roles for other Tolls in *Drosophila* have been elucidated more recently, including functions during wound healing, tube morphogenesis, as well as during axis elongation (Carvalho et al., 2014; Kolesnikov and Beckendorf, 2007; Pare et al., 2014), with each of these roles impinging on actomyosin activity. The role in wound healing requires the standard downstream cascade, including MyD88 as well as the downstream IRAK kinases Tube and Pelle (fly IRAK4), whereas their requirement during tube formation and axis elongation is unresolved.

In particular the role during axis extension, termed germband extension in *Drosophila*, deserves further attention. Here, Toll receptors that in the early fly embryo are expressed in a striped pattern downstream of anterior-posterior patterning machinery localise to apical junctions in the epithelial cells, and mis-expression studies show that they can drive accumulation of actomyosin at the level of adherens junctions. Furthermore, this role appears to depend on heterophilic Toll-Toll receptor interactions, which were confirmed in a heterotypic expression system (Pare et al., 2014). The potential for TLR-TLR interaction was already suggested for the original Toll (Keith and Gay, 1990), but is thus far better characterised in flies than in mammals (Anthoney et al., 2018; Ward et al., 2015). Many family members of the wider LRR-type group are engaged in homophilic and heterophilic interactions (Özkan et al., 2013), supporting the notion that such interactions could be a key part of TLR function. Such function would also place TLRs in a common group with other homophilic/heterophilic cell surface receptors such as Cadherins, Nectins and Crumbs: their levels and ability to engage in trans with receptors on neighbouring cells have important roles in patterning actomyosin activity within the apical-junctional region of epithelial cells (Chang et al., 2011; Thompson et al., 2013).

Junctional patterning of actomyosin activity plays key roles during morphogenesis of epithelial tissues, allowing for defined changes in the shape of the apical domain of epithelial cells (Harris, 2018). Furthermore, junctional tension due to actomyosin contractility is clearly important for the assembly and maintenance of both adherens junctions as well as tight junctions (Citi, 2019; Itoh et al., 2012; Terry et al., 2011).

As a type I-transmembrane protein, a TLR’s biosynthesis involves trafficking to the correct cellular membrane destination. Fly Tolls, where this has been analysed, appear to be localised near apical junctions in the embryonic epidermis (Pare et al., 2014; Towb et al., 1998). The steady-state distribution for mammalian TLRs depending on the TLR type varies between plasma membrane and endosomal localisation, where this has been analysed (Tan and Kagan, 2016). Similar to flies, a number of human TLRs are expressed in epithelial tissues (http://www.proteinatlas.org) and some of these human TLRs show a polarised distribution to either apical or basolateral plasma membrane domains in the lung (Ioannidis et al., 2013) and gut (Price et al., 2018). This polarisation, at least for innate immunity-related functions, most likely reflects an adaptation to where the recognised target PAMPs are most likely to be encountered.

While TLRs have been linked to alterations in epithelial barrier function in the presence of bacteria (Al-Sadi et al., 2021; Guo et al., 2013; Kuo et al., 2013; Ragupathy et al., 2014; Ruffner et al., 2019), little is known about whether they share a physiological role similar to fly Tolls in establishing and maintaining tissue integrity. Here we set out to uncover whether TLRs and downstream components play a non-immune role in mammalian epithelial cells, using differentiated Caco-2 cells as an epithelial intestinal model system. Caco-2 cells show a polarised and junctional localisation of a subset of TLRs. Using inhibitors of the downstream pathway kinase IRAK4 as well as RNA interference against pathway components including TLRs, we show that at steady-state a baseline level of pathway activation is detected, and that absence of this signalling leads to disruption at the tight junction level including loss of junctional tension and a strong reduction in trans-epithelial electrical resistance and thus epithelial barrier function. We suggest that baseline TLR signalling, downstream of TLR-TLR interaction between neighbouring cells in the apical junctional domain, serves a homeostatic role in re-enforcing junctional tension downstream of actomyosin at tight junctions, thereby promoting epithelial tightness.

## Results

### Toll-like receptors are localised to apical junctions in differentiated Caco-2 cells concomitant with steady-state pathway activation

Caco-2 cells grown on transwell filters differentiate into highly polarised enterocyte-like cells, with well-defined junctional regions, dense apical microvilli and well-separated apical and basolateral domains. At 1-3 weeks post-confluence TLR1, TLR2, TLR4 and TLR6, all TLRs previously described to localise to the plasma membrane, were all expressed in Caco-2 cells, and all localised to the apical plasma membrane as well as apical-junctional regions (Fig. 1A-B’’’ and Supplemental Fig. S1). At junctional regions the TLRs colocalised with the junctional enrichment of F-actin (Fig. 1A’’’) as well as tight junction components (Fig. 1B’’’). Similarly, the MyD88 adaptor protein that links the TLRs at the plasma membrane to the downstream IRAKs (Fig. 1F, schematic) was also localised apically and junctionally (Fig. 1C-C’’’). Biochemical analysis revealed that TLR2, TLR4, and TLR6 levels remained constant over the time in culture, whereas TLR1 levels dropped with differentiation (Fig. 1D).

**Figure 1.**
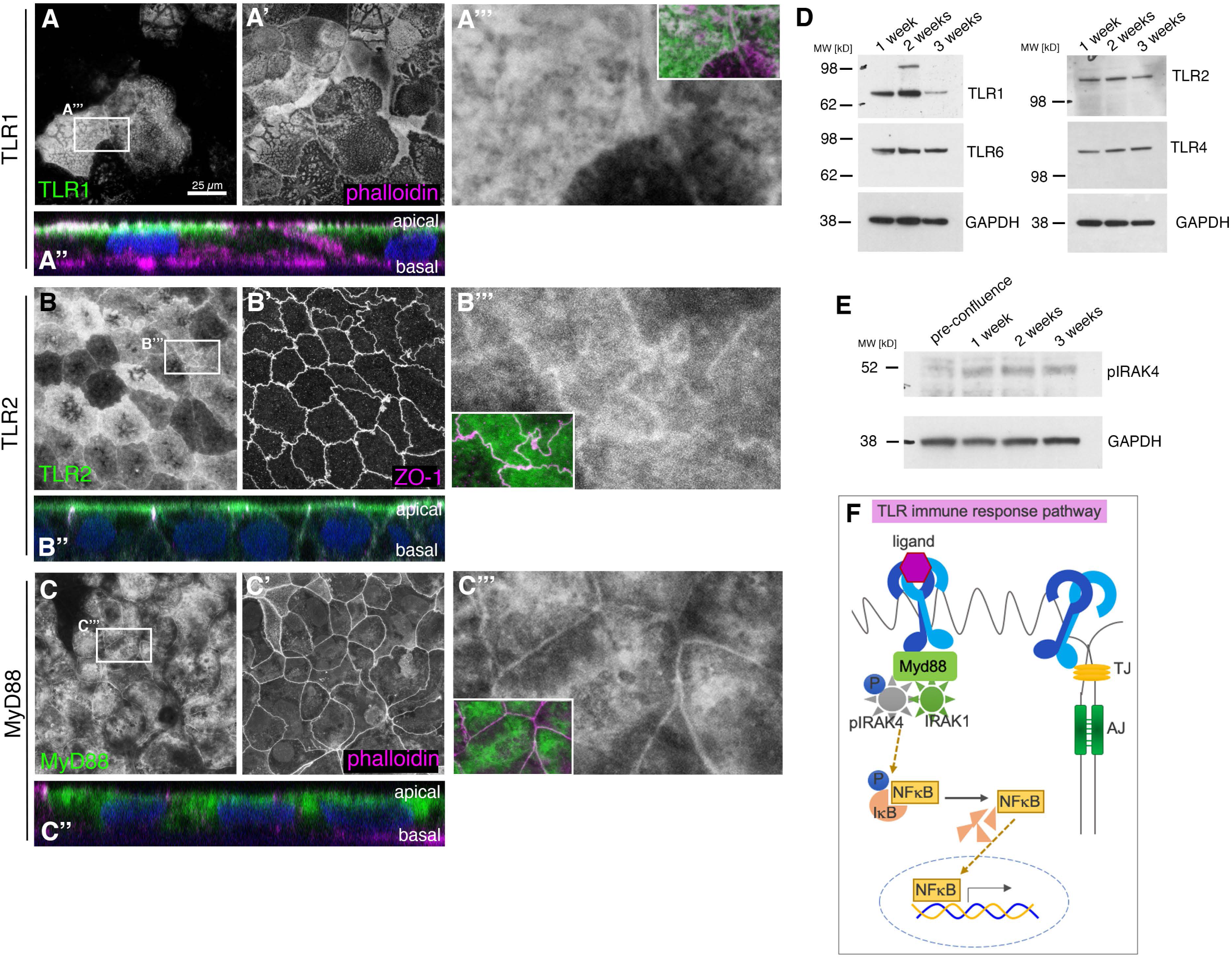
TLRs are constitutively present and localised in a polarised position in epithelial Caco-2 cells. Localisation of TLR1 (**A-A’’’**), TLR2 (**B-B’’’**) and the downstream scaffold protein MyD88 (**C-C’’’**) in epithelial Caco-2 cells. Z-projections of apical-most confocal sections are shown. Antibody labelling against TLR1, 2 and MyD88 is in green, phalloidin and ZO-1 in magenta are used to label cell boundaries. **A’’**, **B’’** and **C’’** show an apical-basal cross-section of the epithelium (nuclei stained with DAPI in blue), apical is up. **A’’’**, **B’’’** and **C’’’** show magnifications of the channels for TLR1, 2 or MyD88, respectively, the insets show the overlay. **D** TLR1, TLR2, TLR4 and TLR 6 are all detected in confluent Caco-2 cells during maturation over 3 weeks by Western Blotting. Note that TLR1 levels consistently drop in fully matured cells. **E** Levels of phospho-IRAK4 (pIRAK4) increase with maturation of Caco-2 cells in culture, indicating steady-state pathway activation and increased signalling with more mature junctions. **F** In innate immunity signalling, TLR binding of an immuno-active ligand triggers assembly of the Myddosome containing pIRAK4, IRAK1 and MyD88, leading to IκB phosphorylation, dissociation from NFκB that is now free to enter the nucleus and affect transcription. A constitutive, non-immune role of the TLR pathway would not require a ligand but might go through the same pathway. See also Supplemental Figure S1.

The presence of TLRs on the epithelial surface, in the absence of any infection, appeared to trigger activation of the downstream pathway (Fig. 1F), as we could detect phosphorylated IRAK4 (p-IRAK4) in these cells, and pIRAK4 levels increased with the cells reaching confluence and differentiating (Fig. 1E). The presence of low-level p-IRAK4 is consistent with findings in other lysates of kidney and immune cell lines (Vollmer et al., 2017). This base-line TLR signalling via pIRAK4 did not seem to involve nuclear translocation of NFκB, as we could not detect NFκB in the nucleus either in control Caco-2 monolayers compared to positive control (Supplemental Fig. S1 C-F).

Thus, intestinal epithelial Caco-2 cells showed a robust and polarised expression of TLRs as well as a baseline activation of the TLR pathway in the absence of infection and thus absence of immune triggers of the pathway.

### IRAK4 inhibition leads to reversible loss of epithelial integrity at tight junctions

In order to test whether TLR pathway activation leading to IRAK4 phosphorylation was important for epithelial function, we blocked IRAK4 function using two commercially available inhibitors of IRAK4, PF06650833 (PF; (Lee et al., 2017)) and AS244697 (AS; (Kondo et al., 2014)) in a range from 1 – 10 µM (Fig. 2A and B). TLR signalling had previously been linked to regulation of epithelial tightness, though usually under infection paradigms (Clarke et al., 2011; Kuo et al., 2013; Ragupathy et al., 2014; Ruffner et al., 2019). We therefore decided to initially focus on the effect of IRAK4 inhibition on epithelial barrier function and tightness.

**Figure 2.**
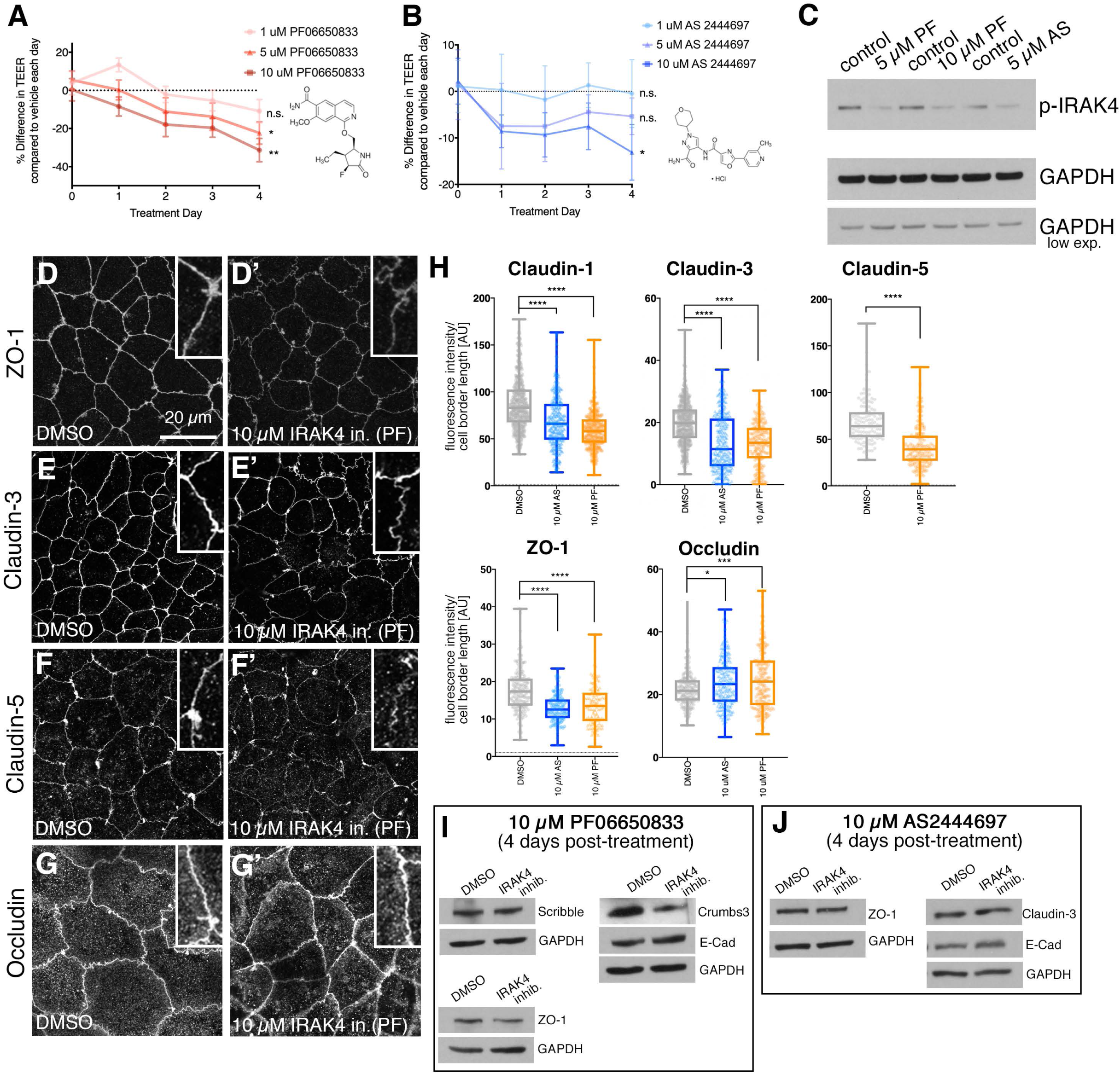
IRAK4 inhibition affects tight junction integrity and barrier function. Treatment of 3-week post-confluent Caco-2 cell monolayers with either PF06650888 (**A**; PF) or AS 2444697 (**B**; AS) IRAK4 inhibitor induces a dose-dependent reduction in TEER. Chemical structures of each inhibitor are indicated. Shown are SEM of n= 6 (DMSO), 6 (AS), 4 (PF) separate transwells. Statistical significance at 4 day time point was determined by one-way ANOVA with Tukey’s multiple comparison test. Note that both AS and PF are compared to same DMSO controls, run contemporaneously. **C** Treatment with 5µM and 10µM PF inhibitor or 5µM AS inhibitor for IRAK4 leads to a reduction in p-IRAK4. **D-H** Effect of PF and AS inhibitor-treatment on junctional components. **D-G’** Z-projections of confocal sections covering the apical-lateral junctional area are shown for cells treated with PF-inhinitor, with quantification for both PF- and AS-inhibitor treated cells shown in **H**. ZO-1 (**D, D’**), Claudin-3 (**E, E’**) and Claudin-5 (**F**, **F’**) localisation at cell-cell junctions is reduced upon IRAK4 inhibition by with 10µM PF, whereas Occludin is slightly increased (**G, G’**). **H** Quantification of fluorescence intensity changes at cell-cell junctions in control (DMSO), 10µM AS- and 10µM PF treated Caco-2 epithelial cell layers. Total junctions analysed from 3 separate images were: Claudin-1: 547 for DMSO, 413 for 10µM AS, 368 for 10µM PF; Claudin-3: 602 for DMSO, 358 for 10µM AS, 211 for 10µM PF; Claudin-5: 120 for DMSO, 192 for 10µM PF; ZO-1: 194 for DMSO, 127 for 10µM AS, 142 for 10µM PF; Occludin: 221 for DMSO, 262 for 10µM AS, 204 for 10µM PF. Box-and-whisker plots in this and all subsequent figures show mean, 25^th^ and 75^th^ percentile, with extreme data points indicated by whiskers. Statistical significance was determined by unpaired Student’s T test or one-way ANOVA with Dunnett’s multiple comparison test when comparing two or more drug treatments (**** = p<0.00001; *** = p<0.0001; * = p<0.01). **I, J** Analysis of protein levels for control (DMSO)- and 10µM PF-treated (**I**) or 10µM AS-treated (**J**) Caco-2 cell monolayers. See also Supplemental Figure S2.

Epithelial monolayers of Caco-2 cells treated with either inhibitor showed a dose-dependent decrease in trans-epithelial electrical resistance (TEER), a measure of tight junction integrity and epithelial tightness, in a reversible fashion (Fig. 2 A, B and Supplemental Fig. S2 A, B). The inhibition was targeting IRAK4, as levels of pIRAK4 decreased in a dose-dependent manner (Fig. 2C). We analysed various components of tight junctions in order to assess what aspect of barrier function had been compromised by the IRAK4 inhibition. The cytoplasmic adaptor of tight junctions, ZO-1, showed a marked decrease in localisation to tight junctions upon IRAK4 inhibition with either PF or AS inhibitor (Fig. 2 D, D’ and H; reduction of 27.2%/ 22.1%, respectively), whereas intensity of the transmembrane protein Occludin slightly increased (Fig. 2 G, G’ and H; increase of 10.1%/13.7% respectively). Claudins form another family of tight junction proteins, characterised by four transmembrane domains. Claudin-1, −3 and −5 all showed a marked loss from tight junctions upon inhibitor treatment (Fig. 2E-F’ and and H, and Supplemental Fig. S2C, C’; reduction of 20.5%/30.0% for Claudin-1, 42.2%/31.8% for Claudin-3 and 38.3% (PF) for Claudin-5). E-Cadherin remained localised to adherens junctions, but the extent of lateral spot adherens junctions basal to the adherens junction belt seemed reduced (Supplemental Fig. S2 D-F). The changes at tight and adherens junctions appeared to be caused by relocalisation of proteins away from the junctions rather than degradation and loss of protein, as at the total protein level no marked differences could be detected (Fig. 2 I and J).

These data indicate that loss of the pIRAK4 signal causes a loss of epithelial barrier function in Caco-2 monolayers, suggesting that the above observed baseline activation of the pathway at steady state could serve to re-enforce the barrier, even in the absence of an immune response.

### Knock-down of IRAK4 leads to loss of epithelial integrity at tight junctions

In order to further confirm the effects observed under IRAK4 inhibition, we decided to also reduce IRAK4 levels by siRNA. As 3-week post-confluent Caco-2 monolayers could not efficiently be transfected, we established a regime of siRNA treatment at confluent seeding and compared epithelial tightness and junctional composition at 1-week post-seeding when the control monolayer had reached a minimum resistance of 600 Ω/cm^2^. Caco-2 cell monolayers treated with siRNA directed against IRAK4 showed a marked decrease in total IRAK4 (Fig. 3A). The monolayers also displayed a consistent and statistically significant reduction in TEER (Fig. 3B). When we analysed tight junction components, ZO-1 showed a clear reduction in fluorescence intensity at cell boundaries (Fig. 3C-E; reduction of 44.2%), similar to what was observed when Caco-2 monolayers were treated with either AS or PF inhibitor. Furthermore, in line with our observations under inhibitor treatment, Claudin-3 and Claudin-5 at junctions were reduced (Fig. 3 F-H for Claudin-3, 25.5% reduction; Fig. 3 I-K for Claudin-5, 37.1% reduction).

**Figure 3.**
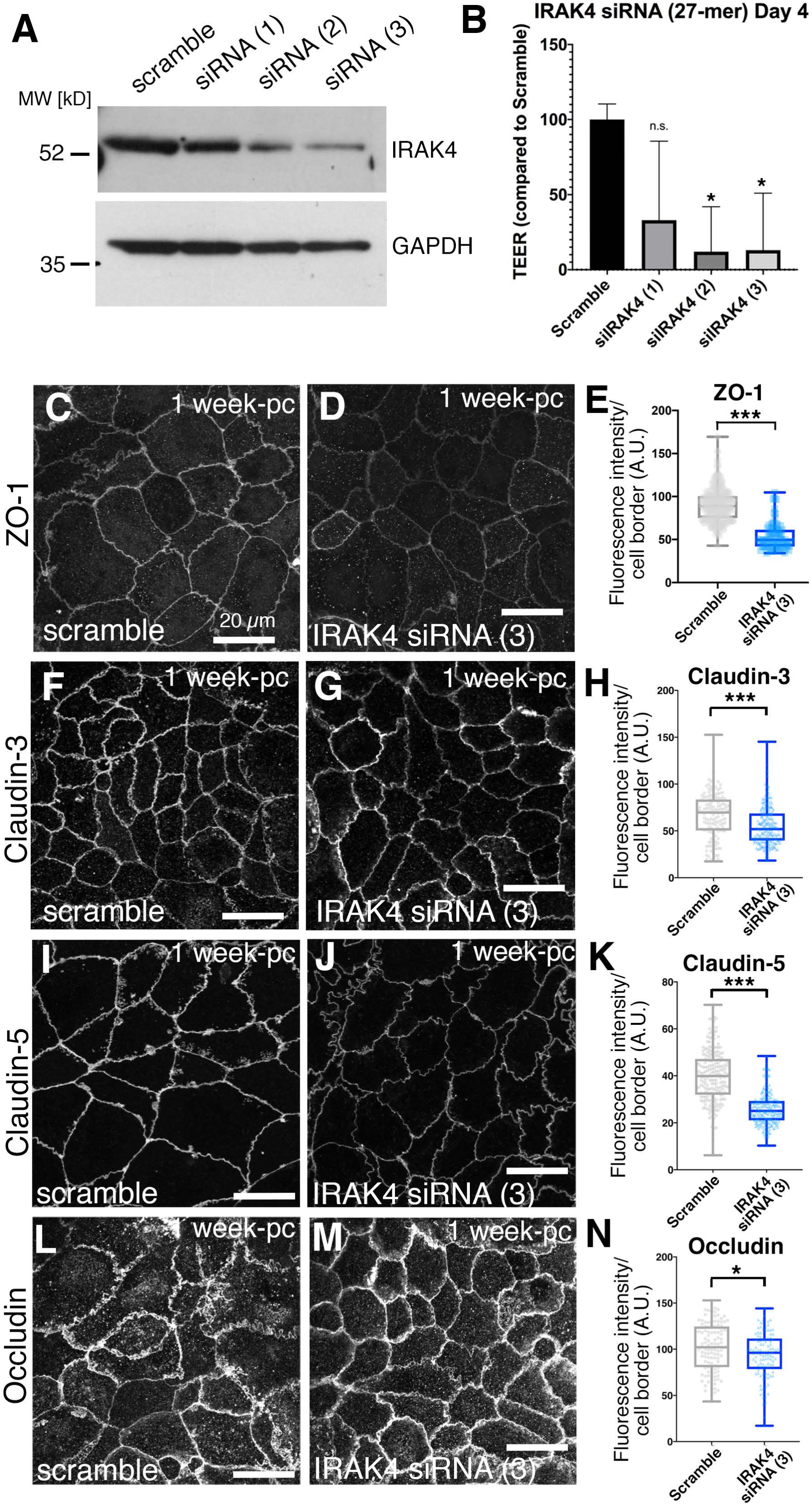
TLR interference phenocopies IRAK4 inhibition and reduction. **A** siRNA treatment of Caco-2 monolayers with three different siRNAs targeting IRAK4 compared to a scrambled control leads to various levels of reduction in IRAK4 levels. **B** In siRNA-treated cells with reduced IRAK4 levels (siRNAs 2 & 3), TEER at day 4 of siRNA treatment is strongly reduced compared to the scrambled control. n=3 transwells were analysed. Error bars indicate SD. Significance was determined by One-way ANOVA and Dunnett’s multiple comparison test. * = p<0.05. **C-E** In Caco-2 monolayers treated with siRNA(3) against IRAK4, ZO-1 levels at junctions are reduced by 44.2% compared to scramble control. **E** n= 250 junctions were analysed for the scramble control and n= 209 for IRAK4 siRNA (3). Statistical significance was determined using unpaired t-test as p<0.00001 (****). **F-H** In Caco-2 monolayers treated with siRNA(3) against IRAK4, Claudin-3 levels at junctions are reduced by 25.5% compared to scramble control. **H** n= 141 junctions were analysed for the scramble control and n= 258 for IRAK4 siRNA (3). Statistical significance was determined using unpaired t-test as p<0.00001 (****). **I-K** In Caco-2 monolayers treated with siRNA(3) against IRAK4, Claudin-5 levels at junctions are reduced by 37.1% compared to scramble control. **K** n= 202 junctions were analysed for the scramble control and n= 227 for IRAK4 siRNA (3). Statistical significance was determined using unpaired t-test as p<0.0001 (***). **L-N** In Caco-2 monolayers treated with siRNA(3) against IRAK4, Occludin levels at junctions are slightly reduced by 5.9% compared to scramble control. **K** n= 111 junctions were analysed for the scramble control and n=110 for IRAK4 siRNA (3). Statistical significance was determined using unpaired t-test as p=0.0147 (*). In **C, D, F, G, I, J, L, M** Z-projections of confocal sections covering the apical-lateral junctional area are shown.

These results confirm the findings obtained with chemical inhibition of IRAK4 described above and support the idea that steady-state TLR signalling via p-IRAK4 in epithelial monolayers plays a role in the re-enforcement of epithelial barrier tightness. Treatment with IRAK4 inhibitor affected monolayers more homogeneously than siRNA and could be applied to more mature monolayers. We therefore performed our further analyses using both chemical inhibition of IRAK4 as well as siRNA induced reduction in protein levels, as these complement each other in terms of efficacy and timepoint of treatment.

### Loss of epithelial integrity downstream of IRAK4 inhibition is due to loss of junctional tension

A key aspect of the establishment and maintenance of tight junctions is tension exerted onto these junctions by actomyosin activity (Fig. 4A) (Itoh et al., 2012; Miyake et al., 2006; Terry et al., 2011). We therefore analysed non-muscle myosin IIA (NMIIA) as well as junctional components involved in anchoring of actomyosin to junctions under control and IRAK4-inhibitor treatment conditions. NMIIA in the apical region of Caco-2 cells at 3-weeks post-confluence was organised into striated patterns, suggesting a mini-sarcomere-like arrangement of actomyosin within this region (Fig. 4B, B’). This striated pattern also demarcated the cell-cell junction region of the cells (Fig. 4C’’). Under IRAK4 inhibitor treatment (PF), NMIIA was less well organised (Fig. 4D-E’’), with foci of myosin across the apical surface as well as near junctions, but many fewer clear striations (Fig. 4E-E’’). NMIIA labelling also extended further basally along the lateral junctions compared to the control, where the highest intensity of NMIIA labelling was confined to the apical and apical-junctional region (compare Fig. 4B’ and D’).

**Figure 4.**
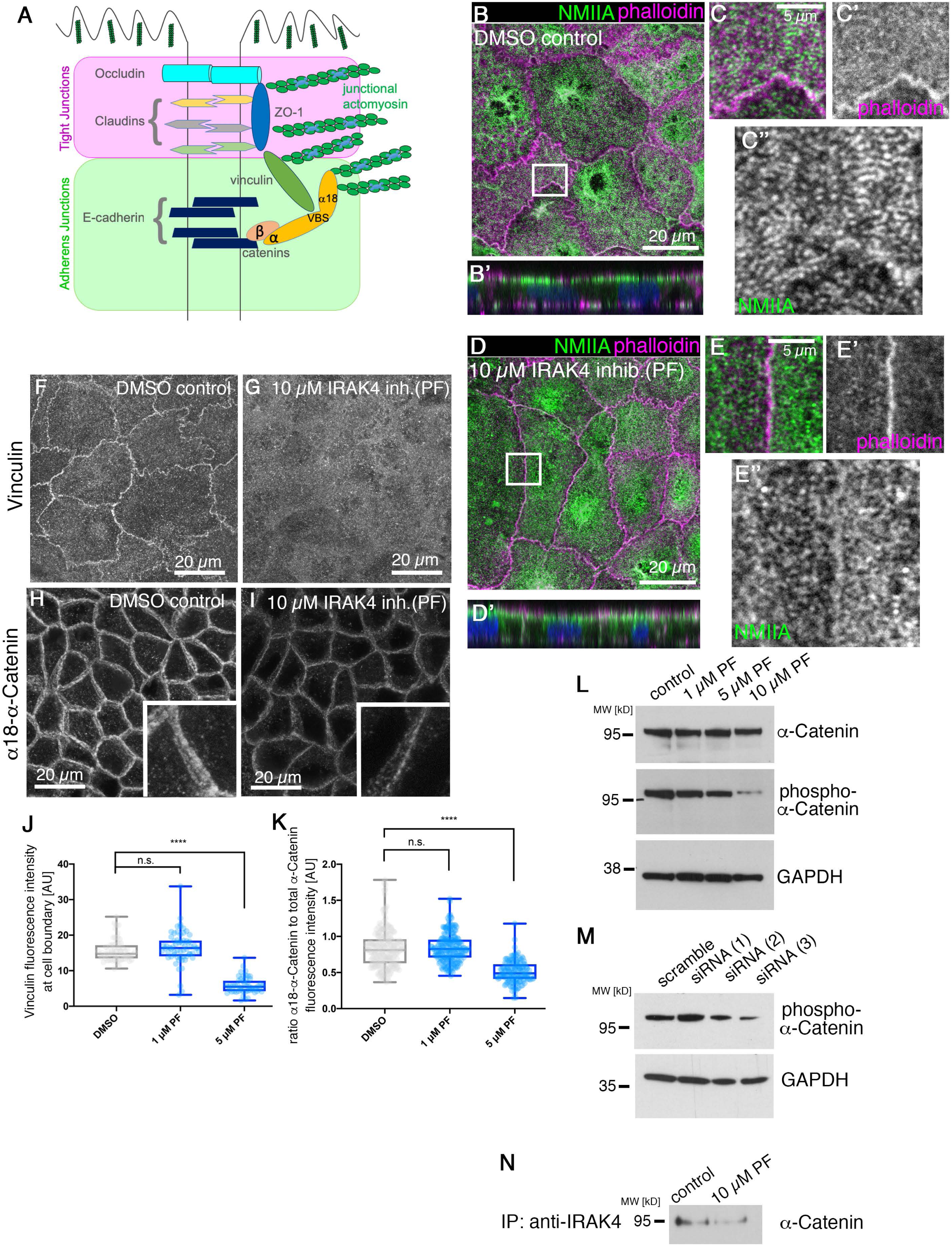
IRAK4 inhibition leads to loss of epithelial tension at tight junctions. **A** Schematic illustrating junctional transmembrane and cytoplasmic components of adherens and tight junctions. **B-C’’** Apical NMIIA in differentiated Caco-2 cell monolayers shows an intricate striated pattern across the free apical surface (**C’’)** and along cell-cell junctions (**B-B’’**), whereas upon 10µM PF treatment (**D-E’’**) NMIIA apical localisation is much more diffuse und unstructured (**E’**’). NMIIA is in green and phalloidin to label F-actin is in magenta (**B, B’, C, D, D’, E**). Z-projections of confocal sections covering the apical surface and junctional area are shown. **F-I** The mechanosensitive cytoplasmic junctional proteins Vinculin (**F**) and α-Catenin (**H**) are reduced upon treatment with 10 µM PF inhibitor to IRAK4 (**G, I**). α-Catenin (**H,I**) is revealed using an antibody against the open and thus stretched conformation of α-Catenin (α18). Z-projections of confocal sections covering the apical-lateral junctional area are shown. Insets show higher magnifications of individual cell’s junctional areas. **J, K** Quantification of changes to junctional Vinculin (**J**) and α-Catenin (**K**) fluorescence intensity in control and IRAK4-inhibitor (PF) treated differentiated Caco-2 cells. α-Catenin intensity is expressed as the ratio between α-Catenin under tension revealed by α18 and total α-Catenin. For vinculin, n= 62 junctions were analysed for the scramble control, n= 63 for 1µM PF treatment and n= 52 for 5 µM PF treatment. For α-Catenin/α18-α-Catenin, n= 196 junctions were analysed for the scramble control, n= 203 for 1µM PF treatment and n= 143 for 5 µM PF treatment. Statistical significance was determined using one-way ANOVA with Dunnett’s multiple comparison test as either non significant (n.s.) or p<0.00001 (****). **L** Serine-phosphorylation of α-Catenin compared to total α-Catenin is reduced in a dose-dependent manner with PF-inhibitor treatment in Caco-2 cell monolayers. **M** siRNA treatment against IRAK4 of Caco-2 cell monolayers also leads to a reduction in phospho-α-Catenin, shown is treatment with the same 3 siRNAs as shown in Figure 3. **N** α-Catenin can be co-immunoprecipitated with IRAK4, and the interaction is reduced upon PF-inhibitor treatment. See also Supplemental Figure S3.

Both adherens junctions and tight junctions are linked to and regulated by the junctional actomyosin cytoskeleton (Citi, 2019). The cytoplasmic adaptor ZO-1 binds to Claudins and Occludin but also contains an actin-binding domain (Fig. 4A)(Furuse et al., 1994; Itoh et al., 1999a; Itoh et al., 1999b). Furthermore, ZO-1 has also been shown to be able to bind directly to α-Catenin (Maiers et al., 2013), and α-Catenin in this capacity can influence tight junction assembly and integrity. α-Catenin is also a key mechanosensitive component of adherens junctions (Huveneers and de Rooij, 2013). A further component in contact with both adherens and tight junctions is the protein Vinculin that also acts as a mechano-sensor, requiring activation and interaction with actin filaments to unfold (Bakolitsa et al., 2004; Janssen et al., 2006). We therefore used antibodies directed against Vinculin, against total α-Catenin as well as an antibody, called α18, directed against the form of α-Catenin that is under tension (Yonemura et al., 2010). Vinculin in control (DMSO-treated) cells colocalised with apical junctions (Fig. 4F), whereas upon treatment with IRAK4-inhibitor (PF) the protein was diffuse and not concentrated at apical junctions (Fig. 4G and J), though protein levels did not appear affected (Supplemental Fig. 4F). When analysing α-Catenin, while total α-Catenin intensity at junctions was slightly reduced upon IRAK4 inhibition (Supplemental Fig. 4C), α18 labelling at junctions was further reduced as a proportion of total α-Catenin, suggesting a loss of tension (Fig. 4 H, I, K). Only the stretched form of α-Catenin that is recognised by the α18 antibody can be bound by Vinculin (Seddiki et al., 2018), therefore the change in α18-α-Catenin observed could be upstream of the observed change in Vinculin.

Recent data show that phospho-regulation of α-Catenin is another pathway of regulating intercellular adhesion (Escobar et al., 2015). Interestingly, levels of phosphorylated α-Catenin were decreasing in a dose-dependent manner with increasing amounts of IRAK4-inhibitor used (Fig. 4L) and were also reduced when IRAK4 was inhibited using siRNA (Fig. 4M). This suggests that the TLR-IRAK4 pathway could in fact impinge on α-Catenin as a target. When IRAK4 was immuno-precipitated from Caco-2 monolayer cell lysates, we could detect α-Catenin being co-immunoprecipitated (Fig. 4N), suggesting that IRAK4 and α-Catenin might co-exist in a complex. IRAK4 inhibition reduced the amount of α-Catenin co-immunoprecipitated by IRAK4 (Fig. 4N).

Thus, inhibition of IRAK4 leads to a reduction in actomyosin-generated tension at tight junctions. Because such tension is crucial to maintain junctional integrity, the loss of eithelial tightness upon IRAK4 inhibition is likely a secondary effect of the loss of apical actomyosin tension across the epithelial layer.

### IRAK4 signalling in junction integrity relays a TLR signal

In order to assess whether the effects on epithelial tightness and junctional organisation due to IRAK4 activity in Caco-2 monolayers were downstream of the apical junctional TLRs, we used siRNA directed against a subset of TLRs to study the downstream effects.

When levels of TLR1 or TLR2 were reduced using different sets of siRNAs (Fig. 5 A, B) we observed a significant reduction in TEER (Fig. 5C). Conversely, treatment with a TLR1/2 agonist (CU-T12-9; (Cheng et al., 2015)) using a regime identical to the siRNA treatment timeline, lead to a significant increase in TEER compared to control (Supplemental Fig. S4J). We then analysed junctional components when TLR1 or TLR2 were reduced using siRNA. ZO-1, Occludin, Claudin-3 and Claudin-5 were reduced at cell borders under either treatment (Fig. 5 D-S; ZO-1 reduction: 44.5% [siTLR1], 30.0% [siTLR2]; Occludin reduction: 14.3% [siTLR1], 9.7% [siTLR2]; Claudin-3 reduction: 27.3% [siTLR1], 49.2% [siTLR2]; Claudin-5 reduction: 41.2% [siTLR1], 30.3% [siTLR2]). This was comparable to what we observed when IRAK4 was reduced by siRNA and under inhibitor treatment. Furthermore, we again observed changes in junctional tension when TLR1 and TLR2 were reduced. At 1-week-post confluence actomyosin structures across the apical surface of Caco-2 cells were not yet as elaborated into sarcomere-like assemblies as at 3-weeks-post confluence (Fig. 4B-C’). Nonetheless, there was a clear reduction of junctional NMIIA accumulation when TLR1 or TLR2 were knocked-down by siRNA, similar to IRAK4 knock-down by siRNA (Supplemental Fig. S4 A-E). In addition, siRNA treatments against TLR1, TLR2, Myd88 and IRAK1 all reduced levels of p-α-Catenin in the cells (Fig. 5 T, U), again supporting that changes at tight junctions observed when the TLR-IRAK4 pathway is inhibited or impaired were downstream of changes in junctional tension.

**Figure 5.**
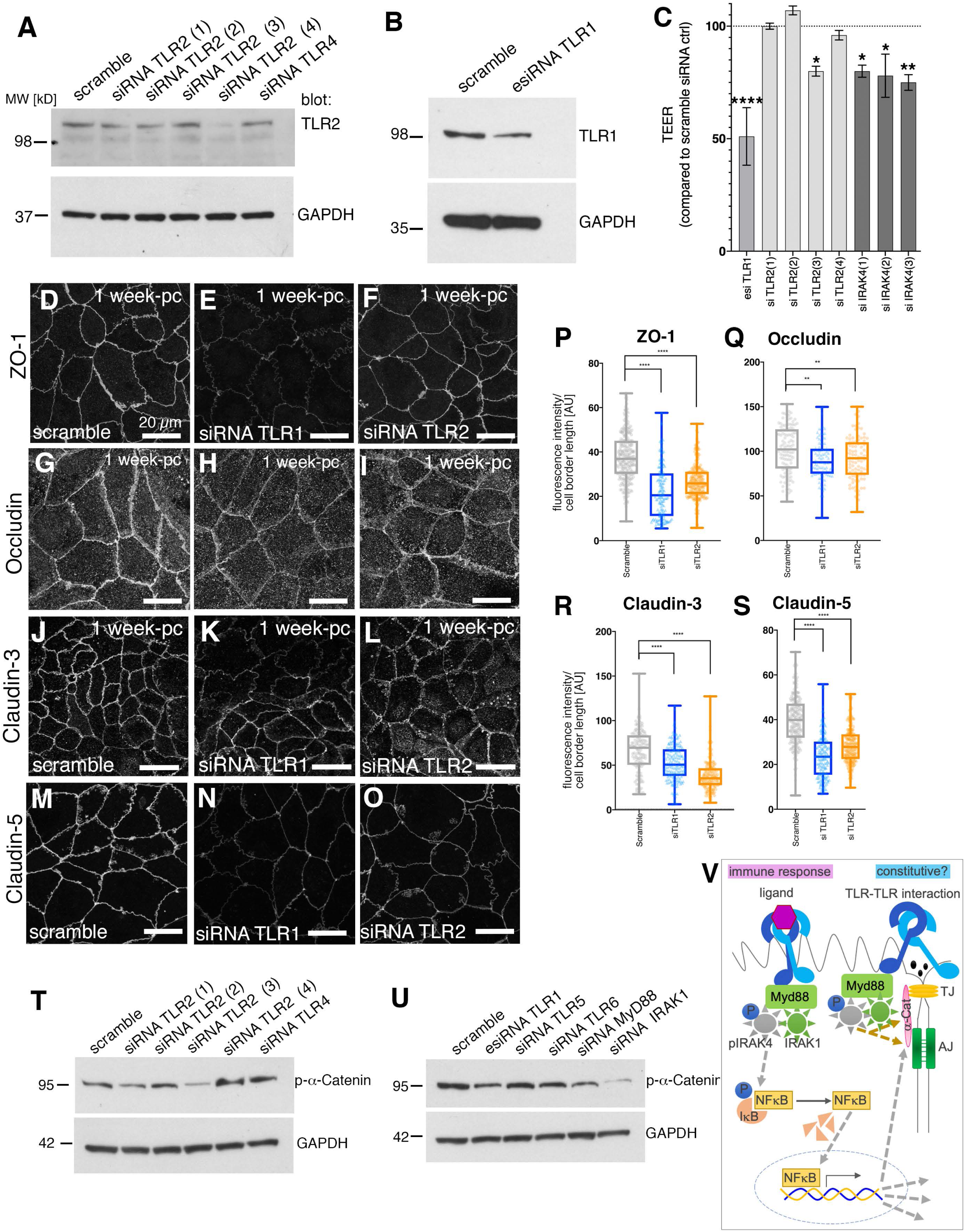
Reduction of TLR levels leads to loss of epithelial tightness and changes at junctions. **A, B** Western blot analysis of protein levels in 1-week post-confluence Caco-2 monolayers upon treatment with siRNAs targeting TLR2 (1-4) or TLR4 as control (**A**) as well as TLR1 (**B**), to demonstrate a reduction in protein level for TLR2 and TLR1. **C** siRNA treatment of Caco-2 monolayers with different siRNAs targeting TLR1, TLR2 or IRAK4 compared to a scramble control leads to reduction in TEER level. n= 5 independent transwells (scramble, siTLR1, siTLR2, siIRAK4) were averaged. Statistical significance was determined by one way ANOVA with Dunnett’s multiple comparison test as * = p<0.01, ** = p<0.001, **** = p<0.00001. **D-O** In siRNA-treated 1-week post-confluence Caco-2 monolayers with reduced TLR1 or TLR2 levels, localisation of junctional components to cell borders is affected. Z-projections of confocal sections covering the apical-lateral junctional area are shown. **D-F** Knock-down of TLR1and TLR2 leads to ZO-1 reduction at junctions of 44.5% and 30.0%, respectively. **G-I** Knock-down of TLR1and TLR2 leads to mild Occludin reduction at junctions of 14.3% and 9.7%, respectively. **J-L** Knock-down of TLR1and TLR2 leads to reduction of Claudin-3 at junctions of 27.3% and 49.2%, respectively. **M-O** Knock-down of TLR1and TLR2 leads to reduction of Claudin-5 at junctions of 41.2% and 30.3%, respectively. **P-S** Quantification of components in scramble control, TLR1 siRNA and TLR2 siRNA treated Caco-2 cells. **P** ZO-1: n= 267 junctions were analysed for the Scramble control and n= 160 for siTLR1 and n=216 for siTLR2. **Q** Occludin: n= 111 junctions were analysed for the scramble control and n= 125 for siTLR1 and n=93 for siTLR2. **R** Claudin-3: n= 141 junctions were analysed for the scramble control and n= 175 for siTLR1 and n=142 for siTLR2. **S** Claudin-5: n= 202 junctions were analysed for the scramble control and n= 168 for siTLR1 and n=187 for siTLR2. Statistical significance was determined using one-way ANOVA with Dunnett’s multiple comparison test as** = p<0.001, **** = p<0.00001. Note that scramble values for Claudin-5 and Occludin are the same as those in Figure 3 as these were run contemporaneously. **T** Analysis of phospho-α-Catenin levels upon treatment with siRNA against TLR2 (1-4) or against TLR4 compared to scramble control. GAPDH is shown as loading control. **U** Analysis of phospho-α-Catenin levels upon treatment with siRNA against TLR1, TLR5, MyD88 orIRAK1 compared to scramble control. GAPDH is shown as loading control. **V** Model of ‘constitutive’ TLR pathway function compared to the immune-response-induced pathway. See also Supplemental Figure S4.

In summary, the above data support a model whereby TLR1 and TLR2, possibly as transcellular heterodimers, are involved in sensing of the epithelial state and through downstream MyD88, IRAK4 and IRAK1 help to mature and maintain junctional tension and hence epithelial tightness.

## Discussion

TLRs comprise a large family of membrane receptors conserved through evolution that fulfil many diverse functions both during development and throughout life. While this family includes both cell surface and endosomal receptors known to respond to different stimuli, the majority of TLR signalling is reported to converge on a common signalling hub, the myddosome, and information is relayed from here to the nucleus. Recent data from *Drosophila*, however, suggests functions that might involve the myddosome but do not effect changes through altering transcription. In agreement with this, our data suggest that a common epithelial role of the upstream part of the pathway might impinge on junctional integrity and hence epithelial barrier function through the modulation of junctional tension.

A link between TLR signalling and epithelial barrier function is well described during infection progression, though stimulation of different TLRs depending on the tissue context can either lead to a decrease or an increase in barrier function. For instance, recent studies suggest that some bacteria during infection of the lung could exploit a transient relaxation of the epithelial barrier downstream of TLR signalling (though through non-canonical downstream components p38 and MAPK) to invade the host (Clarke et al., 2011). This increase in leakiness of the epithelium, characterised by a decrease in expression of the tight junction-associated Claudins as well as increase in SNAIL1 expression (a transcriptional repressor of Claudins), could allow egress of immune cells as well as antimicrobial factors into the lung lumen as part of the host defence. Conversely, differential TLR-mediated control of junction strength in the Peyer’s patches of gut epithelium enhances leakiness to allow localised sampling of the luminal surface by dendritic cells as part of normal immune surveillance (Davies et al., 2010). Interestingly, recent studies of neurobehavioral deficits in TLR2 knockout mice indicated problems at junctions (Hu et al., 2020). TLR2 knock-out mice are viable but display a range of brain-related problems, in agreement with TLR2 being expressed widely in glial cells and neurons in the nervous system (Hayward and Lee, 2014). These mice at 12 months of age displayed blood-brain barrier problems, concomitant with a reduction of protein levels of ZO-1, Occludin and Claudin-5 in this tissue. Several studies of murine gut inflammation support a key role for TLR1 and TLR2 in maintaining barrier integrity. TLR1 knockout mice exhibit decreased proliferation rates in the colonic crypt and impaired recovery of the tissue after colitis induction (Kamdar et al., 2018). TLR2 stimulation had a protective effect on tight junctions in animals, explants and primary human intestinal cells in culture, and both TLR2 and MyD88 knockout mice exhibit an accelerated disruption of the barrier following colitis induction (Cario et al., 2007). Additionally, a recent study demonstrated that TLR1 and TLR2 (but not TLR6) knockout mice display increased intestinal permeability and pathogenic yeast colonisation in a colitis model (Choteau et al., 2017).

How essential or redundant are epithelial barrier function of the TLR pathway and its components? Mutations in some of the genes involved in TLR signalling, such as IRAK4, are compatible with life in both human and mouse in the absence of infection, however in individuals carrying the mutations the innate immune response is reduced and sometimes not effective (Picard et al., 2010). IRAK4 mutants in humans for example show an increased sensitivity to infections, specifically in the upper respiratory tract (Ku et al., 2007). Other TLR mutations, including those TLRs described here (1, 2, 4, 6), have been linked to a broad spectrum of immune disorders and chronic infections, such as colitis in mice and common variable immunodeficiency, asthma, Crohn’s disease, atherosclerosis, tuberculosis, and leprosy in humans (Choteau et al., 2017; Lin et al., 2012; Mortaz et al., 2017). The absence of lethality for many of the mutants suggests redundancy in the system. Indeed, during a phase 1 clinical trial one of the inhibitors used in this study (PF) revealed no evidence of a severely disrupted epithelial barrier (Danto et al., 2019). With regards to IRAK4 inhibition and knock-down, we suspect that the acute effects we observe in either case are masked in the developmental context by such redundancy, hence leading to overall viability.

While this work suggests a role for TLRs in the maintenance of epithelial barrier function, the precise mechanism of TLR/IRAK4 mediated control of junctional components requires further investigation. While our data suggest a complex formation between IRAK4 and α-Catenin, there is no known IRAK4 consensus site on α-Catenin, and recent evidence suggests that the kinase activity of IRAK4 is dispensable for the function of the myddosome (De Nardo et al., 2018). Furthermore, it is not yet clear how IRAK1, the major downstream effector of IRAK4, is involved in the epithelial barrier function, although IRAK1 was recently identified in an affinity biotinylation screen of E-cadherin binding partners (Guo et al., 2014).

A TLR/IRAK4-mediated complex formation that maintains α-Catenin phosphorylation may explain the observed decrease in TEER, which is thought to be primarily a measure of the “pore” pathway mediated by Claudins. α-Catenin is a mechanosensitive link between the circumferential adherens junction and the apicolateral tight junction, and its phosphorylation helps maintain intercellular adhesion in both human cell culture and *Drosophila* (Escobar et al., 2015). In addition, by maintaining the “unfolded” form of α-Catenin to allow recruitment of Vinculin, TLR/IRAK4 signalling may help buffer the tight junction from disruption via mechanical forces (Konishi et al., 2019). Importantly, despite our observations of altered junctional proteins, we do not observe large scale changes in total protein levels under IRAK4 inhibition or siRNA, and, following removal of IRAK4 inhibitors, TEER rapidly recovered to control levels. This suggests a translocation of proteins away from the cell-cell interface rather than protein degradation, although the precise mechanism remains to be determined.

How does the modulation of junctional tension, that we observe when either IRAK4 or TLR1/2 are targeted, fit with other developmental data? In *Drosophila*, one key function for TLRs has emerged during the early morphogenetic process of germband elongation that occurs during gastrulation (Pare et al., 2014). A combinatorial effect of heterophilic TLR-TLR interaction at boundaries of striped expression domains of different TLRs in the embryonic epidermis leads to actomyosin accumulation at such boundaries. Different TLRs were shown to be able to interact heterophilically in a heterologous expression system. We suspect that in the case of epithelial monolayers such as the one analysed here, homo- or heterophilic interactions of TLRs localised to the apical-junctional region relay a steady-state signal to the pathway (Fig. 5V). In Caco-2 cells, our data show, this signal leads to the strengthening of actomyosin interactions with junctions and involves the mechano-sensitive component α-Catenin and its binding partner Vinculin. It will be interesting to address in the future whether the effect fromTLRs onto junctional myosin in *Drosophila* involves the same set of effectors.

## Materials and Methods

### Cell culture, transfection and inhibitor treatment

Caco-2 cells were grown on 24- or 6-well transwell polycarbonate inserts with 0.4μm pore size (Greiner Bio-one) at a cell density of 10^5^ cells/cm^2^. Complete growth medium consisted of DMEM/Glutamax (Gibco 10566016) supplemented with 10% FBS (Gibco), 1% Penicillin/Streptomycin, and 1X MEM non-essential amino acids (Gibco) and was changed every other day. Unless otherwise indicated, cells were grown for 21 days post-confluence to achieve full apicobasal polarization and junction maturation. For transfection studies, cells were reverse-transfected with Lipofectamine 2000 (Invitrogen) with 100ng of plasmid on a 24-well transwell insert in 200μl antibiotic free media as directed by manufacturer. For drug studies, IRAK4 inhibitors (PF-06650833 or AS2444697) were applied as DMSO suspensions to both apical and basal chambers at concentrations of 1, 5, and 10µM, and changed every day for 4 consecutive days. For controls, DMSO was added to media at concentration equal to the amount of DMSO in the highest concentration of drug tested.

### Trans-epithelial electrical resistance (TEER) measurements

TEER measurements were carried out using an EVOM^2^ Voltometer (World Precision Instruments) with STX2 chopstick electrodes. Both samples and calibration media were allowed to acclimatise to room temperature prior to measurements to minimize fluctuations in resistance during cooling. Resistance was calculated using Ohm’s law and is reported as Ωcm^2^. In some instances, data were normalised to matched controls to improve clarity as noted in figure legends. Prior to any treatments, samples below a TEER cut-off of 600 Ωcm^2^ were excluded from the analysis.

For graphs presented in Figure 2, values indicate weighted mean values from 3 independent experiments expressed as percent difference from DMSO controls. To find weighted means, normalised values were averaged by the formula:

(((mean_1_*N_1_)/N_total_)+((mean_2_*N_2_)/N_total_)+((mean_3_*N_3_)/N_total_)).

SEM was determined by propagation of error by the formula:

Sqrt((((SEM_1_^2^)*N_1_)/N_total_)+(((SEM_2_^2^)*N_2_)/N_total_)+(((SEM_3_^2^)*N_3_)/N_total_)). Statistical significance for final timepoint was determined by one-way ANOVA with Dunnett’s multiple comparison test.

### Immunofluorescence and imaging

Cells were washed briefly in PBS and fixed in 4% methanol-free PFA (2% for anti-Claudin antibodies) for 15 minutes at room temperature. They were then washed 3 times and permeabilised in PBS containing 0.1% saponin for 10 minutes, followed by blocking in PBS containing for 0.1% saponin and 1% BSA for 1 hour. Transwell membranes were then excised from plastic supports with a scalpel, and incubated in primary antibodies (see below for full details) overnight at 4°C in blocking buffer. Membranes were then washed 3 times with PBS and incubated with species-matched Alexa Fluor-conjugated secondary antibodies and Phalloidin, followed by counterstaining with DAPI and mounting using Vectashield (Vector Laboratories). For all TLR antibodies and Myd88, signal was enhanced using a Tyramide Superboost kit (Invitrogen). Cells were imaged on a Leica SP8 or Olympus FluoView 1200. Images were processed using Fiji software to adjust colour palette, contrast, and balance; all changes were applied to the entire image and for comparisons the same settings were applied to all images being compared. Z-projections of confocal sections (standard deviation projections) either covering the apical and apical-junctional area or the whole lateral side were analysed (all fluorescence panels shown in figures are standard deviation projections). Fluorescence intensity was quantified using Fiji. Briefly, junctions were traced using the freehand tool with a line thickness matched to the thickest junction within the set of images to be compared (usually 10-15pt). Average signal intensity over junction length/width was measured. For each image, background intensity was sampled and averaged from 4-5 cytoplasmic regions approximately the length of a junction and subtracted from all values within the image. For NMIIA images, ratios were determined by dividing junction intensities by intracellular NMIIA intensity, as determined by averaging 3-4 circular ROIs covering intracellular spaces

Antibodies used:

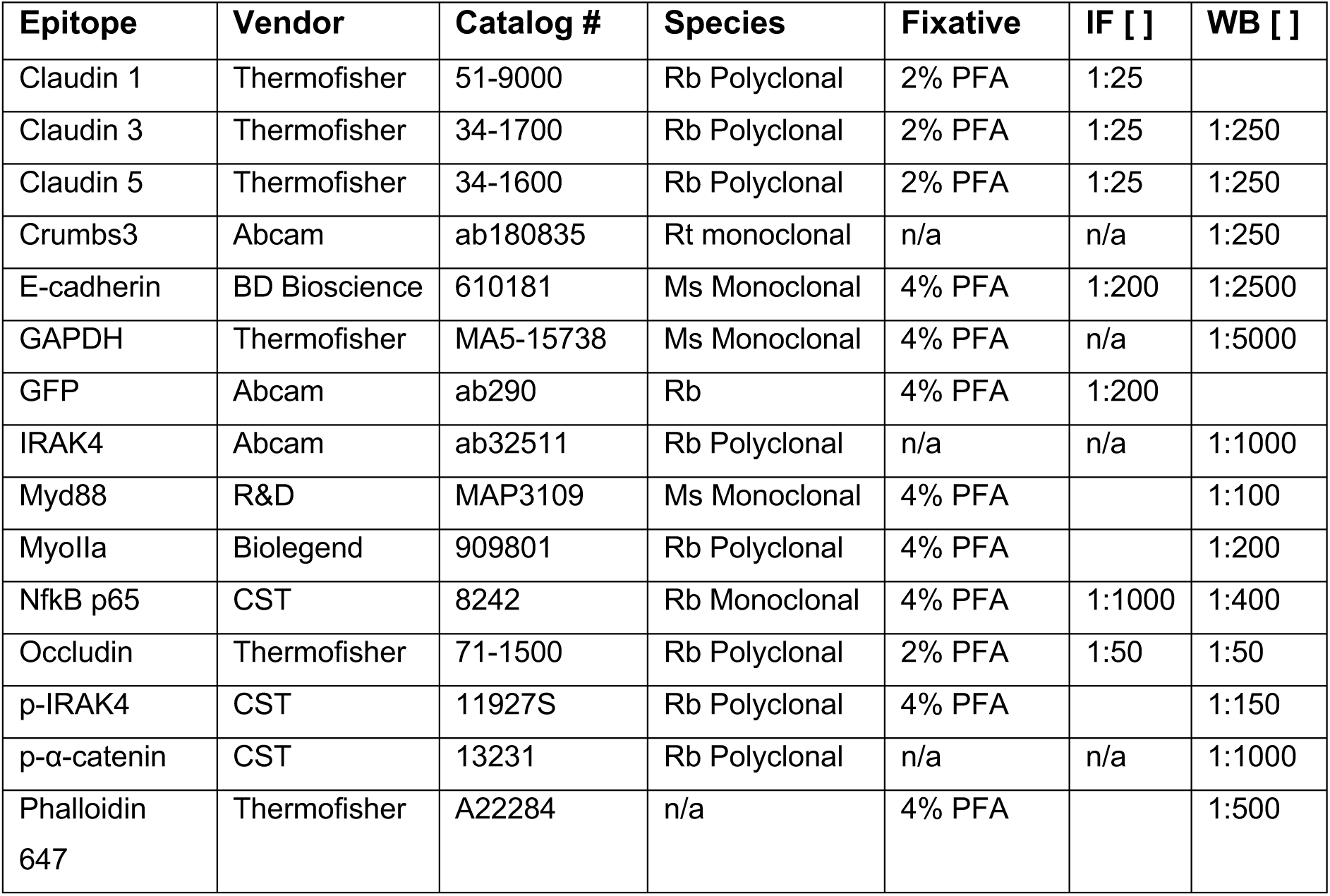
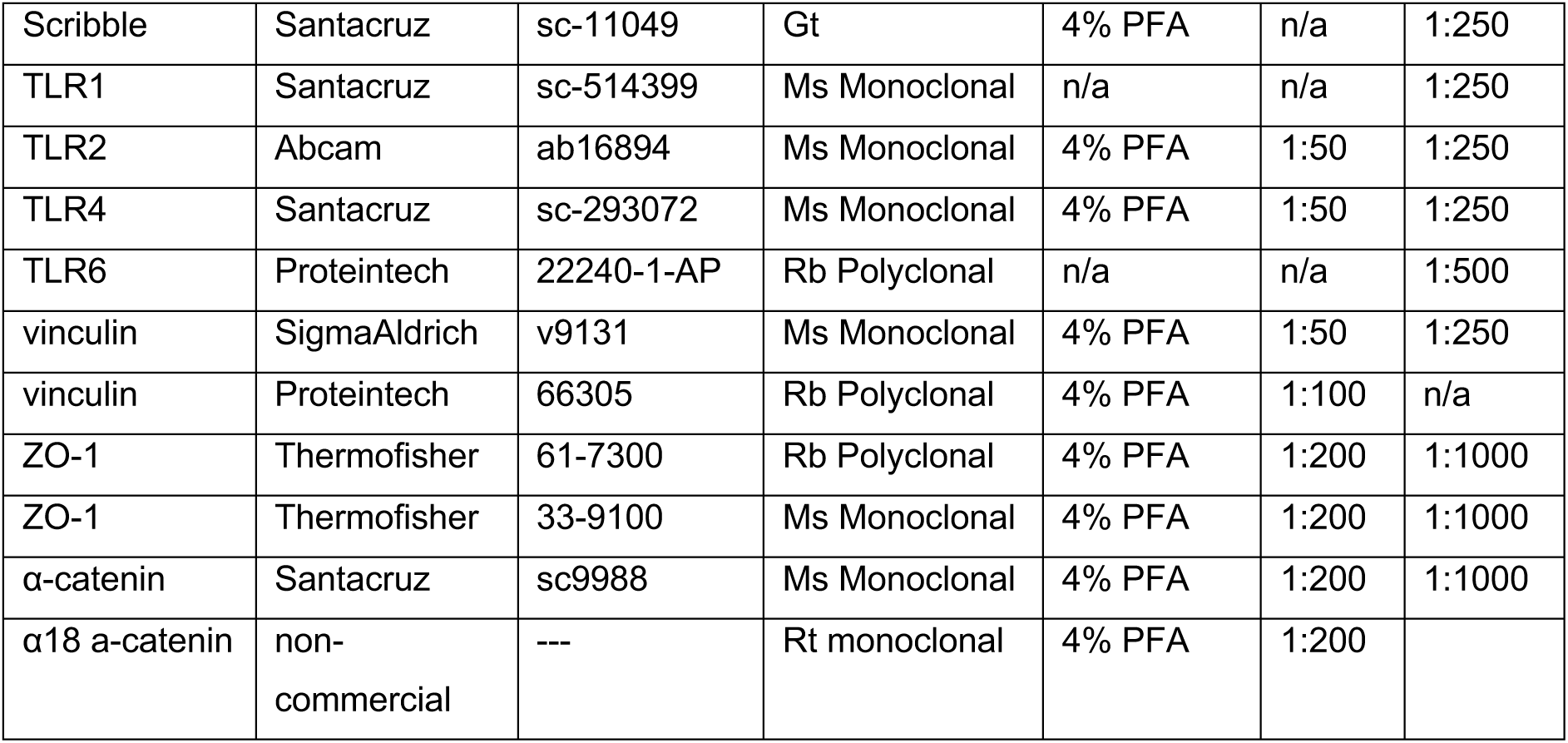

### Constructs for Transfection

Fluorescent TLR1, TLR2, TLR4, TLR6, and Myd88 constructs (13014, 13015, 13016, 13018,13020) were obtained from Addgene.

### Biochemistry and Western Blotting

Samples for Western blot analysis were grown on 6-well transwell polycarbonate inserts with 0.4μm pore size (Greiner Bio-one) at a cell density of 10^5^ cells/cm^2^. All subsequent steps were conducted on ice or at 4°C. For lysis, membranes were washed twice with ice-cold PBS containing cOmplete^TM^ protease inhibitor and phosSTOP^TM^ phosphatase inhibitor cocktails (Roche). Cells were then gently lifted from membranes with cell scrapers (Corning) and centrifuged at 1000 rpm for 5 minutes. Supernatant was removed and cells were resuspended in lysis buffer containing 20mM Tris pH 7.5, 1mM EDTA pH 8.0, 1% Triton X-100, 150mM NaCl, and protease/phosphatase inhibitors (Roche). Samples were passed through a 29g syringe and placed on ice for 1 hour, vortexing for 30 seconds every 15 minutes. Samples were then spun at 10,000rpm for 10 minutes and pellet was discarded. BCA protein assay (Thermofisher) was used to determine concentration on a NanoDrop 2000 (Thermofisher), Samples were then equilibrated to 2µg/µl with lysis buffer, 1x LDS sample Buffer (Invitrogen) and 1mM Dithiothreitol. 40μg of protein were added to each lane and samples were electrophoresed on 4-12% Bis-Tris gels (NuPage, Thermofisher). Gels were then transferred to PVDF membrane, blocked in blocking buffer consisting of PBS, 0.1% Tween 20, and 5% milk (Marvel). Membranes were then probed overnight with primary antibodies (see Table S1) in blocking buffer at 4°C. The following day, membranes were washed 3 times with blocking buffer and incubated for 1 hour in HRP-conjugated secondary antibody. Membranes were then washed 2 times with blocking buffer for 1 hour total, washed briefly with PBST, and incubated with Prime ECL (Invitrogen) for 10 minutes before imaging. Densitrometry was performed with Fiji. For co-immunoprecipitation, lysates from two pooled wells of a 6-well plate were prepared as above, followed by preclearing with 20 µl magnetic beads (Pierce) for 30 minutes at 4°C. After removal of beads, primary antibody was added (1:50) to crude lysates and samples were placed in a rotating rack in a 4°C cold room overnight. Following overnight incubation, 20 µl pre-washed magnetic beads were added to the lysate/antibody mixture. Samples were rotated in the cold room for 1 hour, and pelleted with a magnetic rack. Pellets were washed 5 times with cell lysis buffer and resuspended in 30ul 4x sample buffer (Invitrogen).

### siRNA treatments

The protocol is adapted from a method published on the bio-rad website (https://www.bio-rad.com/webroot/web/pdf/lsr/literature/bulletin_5370.pdf). Briefly, transwells were seeded at high density 10^6^ cells/cm^2^ and reverse transfected with 80 nM siRNA using Lipofectamine 2000. After six hours, media was replaced with fresh antibiotic-free media to remove excess of cells. Overnight confluency was confirmed visually, and after one week knockdown efficacy was confirmed by western blot. siRNAs were selected based on efficacy and included standard siRNAs (TLR2: Dharmacon, LQ-005120-01-0005) as well as 27-mer (IRAK4: Origene SR322049) and esiRNAs (Sigma Mission, TLR1: EHU117701, TLR4: EHU086621, TLR6: EHU022071, IRAK1: EHU093291, Myd88: EHU029771).

### Statistical Analysis

Unpaired Student’s t test was used for single comparisons between normal distributions, while one- or two-way ANOVA with Dunnett’s or Tukey’s method were used for multiple means comparison unless indicated in text. For all statistical measures, the number of images or wells used to generate data are indicated in figure legends. For fluorescence quantification, replicates were taken from at least 3 separate membranes stained and imaged contemporaneously, and Z-stacks were collapsed into a standard deviation projection prior to measurement. Graphs were produced and analysed in Prism. Error bars indicate standard error of the mean.

All box-and-whisker plots show mean, 25^th^ and 75^th^ percentile, with extreme data points indicated by whiskers. N values and statistical tests used are indicated in the relevant figure legends.

---

**Supplemental Figure S1, related to Figure 1.**
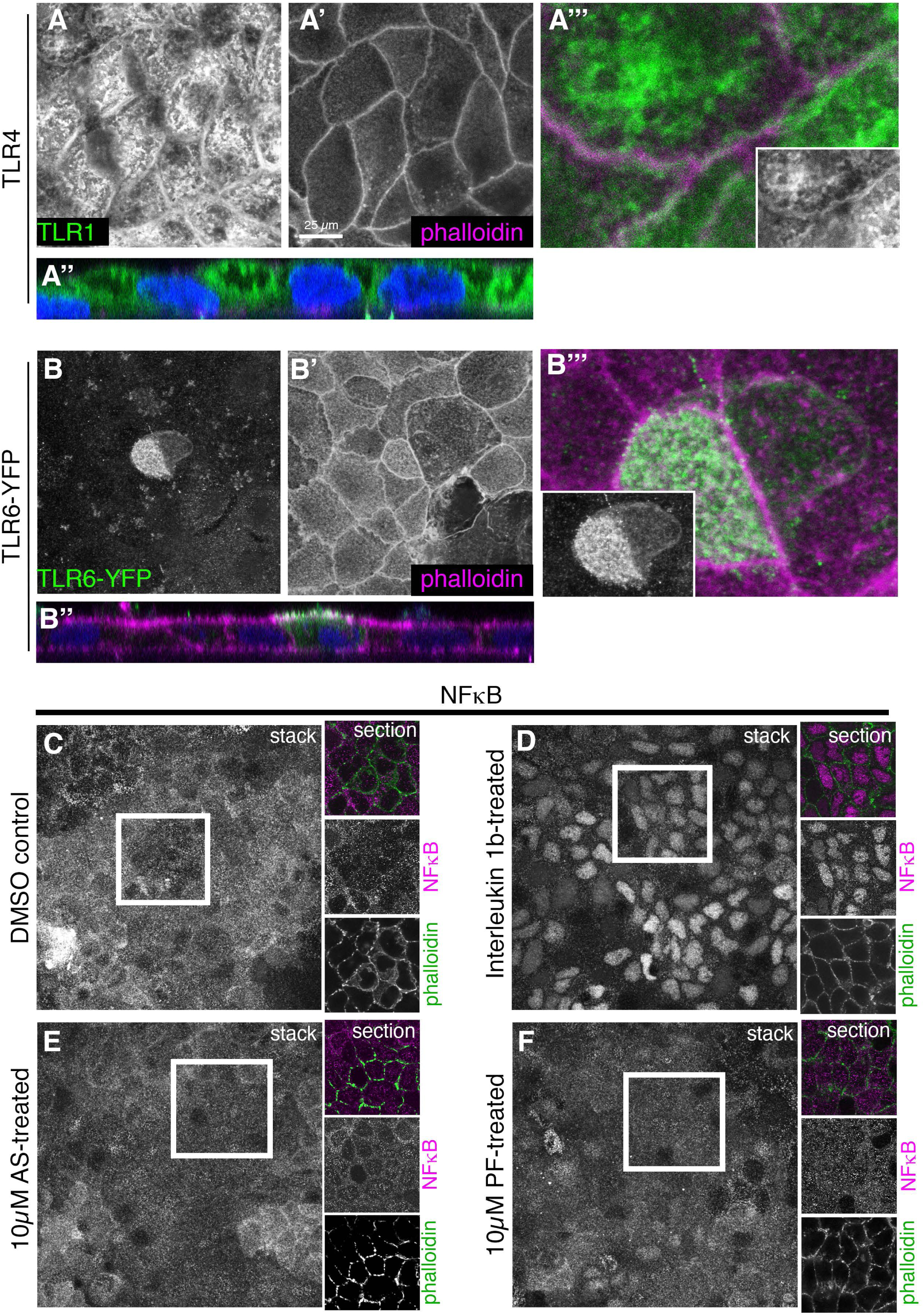
TLRs are constitutively present and localised in a polarised position in epithelial Caco-2 cells. Localisation of TLR4 (**A-A’’’**), TLR6-YFP (**B-B’’’**) in epithelial Caco-2 cells. Z-projections of apical-most confocal sections are shown. Antibody labelling against TLR4 and overexpressed TLR6-YFP are in green, phalloidin to label cell boundaries is in magenta. **A’’**, **B’’** show an apical-basal cross-section of the epithelium (nuclei stained with DAPI in blue), apical is up. **A’’’**, **B’’’** show magnifications of cell boundary regions, the insets show the single channel for TLR4 or TLR6-YFP, respectively. **C-F** Constitutive TLR signaling in Caco-2 monolayers does not involve nuclear NFκB. 3-week post-confluence Caco2 monolayers have little to no nuclear NFκB (**C**) whereas monolayers treated with 10µM Interleukin-1b, a canonical TLR pathway agonist, show strong nuclear NFkB labeling after 30 mins (**D**). Caco2 monolayers treated with IRAK4-inhibitor (see Fig. 2) like the DMSO control show no nuclear NFκB (**E**, **F**). Cell outlines are labelled with phalloidin revealing F-actin.

**Supplemental Figure S2, related to Figure 2.**
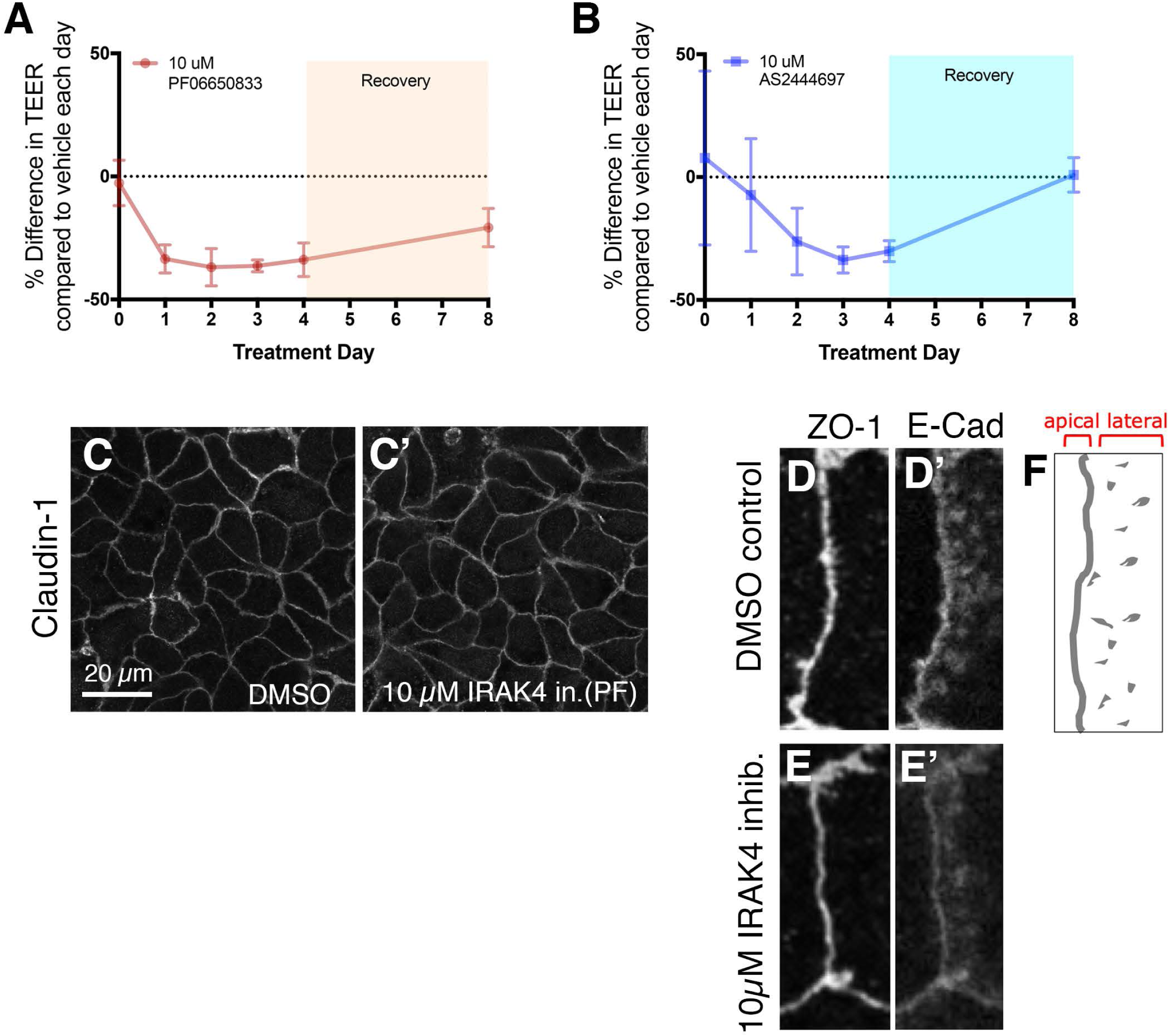
IRAK4 inhibition affects tight junction integrity and barrier function. Treatment of 3-week post-confluent Caco-2 cells with 10µM of either PF06650888 (**A**) or AS 2444697 (**B**) IRAK4 inhibitor induces a dose-dependent reduction in TEER that is reversible upon inhibitor removal (recovery), indicating that the epithelial barrier at tight junctions is re-enforced upon washout of the inhibitor. Shown are SEM of n=3 separate transwells per excperimental condition (DMSO, AS, PF). Statistical significance for the final timepoint was determined by unpaired Student’s t-test. **C-C’** Effect of 10µM PF inhibitor-treatment on junctional components. Z-projections of confocal sections covering the apical-lateral junctional area are shown. The junctional intensity of Claudin-1 is decreased compared to control (DMSO)-treatment. Quantification is in Fig. 2H. **D-F** E-Cadherin at apical junctions appears to change its distribution within lateral spot adherens junctions (see schematic in **F**) upon 10µM IRAK4 inhibitor-treatment (PF). ZO-1 (**D**, **E**) is shown in comparison to E-Cadherin (**D’**, **E’**) and to mark the apical-most end of the lateral sides.

**Supplemental Figure S3, related to Figure 4.**
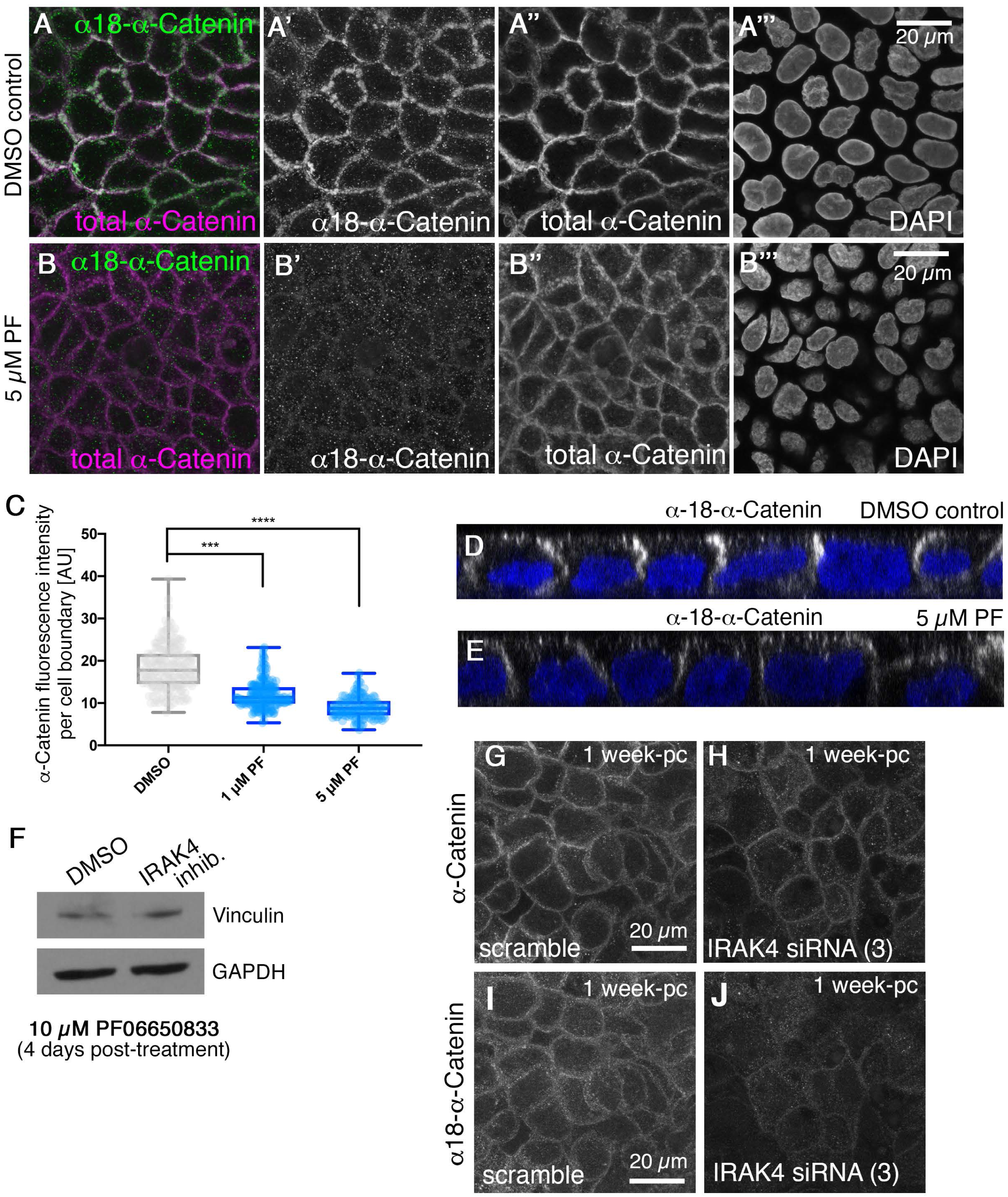
IRAK4 inhibition leads to loss of epithelial tension at tight junctions. **A-B’’’** Immunofluorescence of α18-α-Catenin compared to total α-Catenin (as quantified in Fig. 3K). Z-projections of confocal sections covering the apical-lateral junctional area are shown. **C** Total α-Catenin fluorescence intensity at junctions in control and PF-inhibitor treated Caco-2 cells. Values are extracted from α-Catenin/α18-α-Catenin ratios (Figure 4K) and statistical significance was determined using one-way ANOVA with Dunnett’s multiple comparison test as either p<0.0001 (***) or p<0.00001 (****). **D, E** Cross-sections corresponding to panels **A’** and **B’** above illustrating α18-α-Catenin localisation at lateral junctions in control (DMSO) and 5µM PF-inhibitor treated Caco-2 cell monolayers. **F** Vinculin total protein levels do not change between control and PF-inhibitor-treated Caco-2 cells, as anlaysed by Western blotting. **G, J** In Caco-2 monolayers treated with siRNA(3) (see Figure 3) against IRAK4, both α-Catenin (**G**, **H**) as well as α18-α-Catenin (**I**, **J**) levels at junctions are reduced compared to control, with changes to α18-α-Catenin stronger than to total α-Catenin. Z-projections of confocal sections covering the apical-lateral junctional area are shown.

**Supplemental Figure S4, related to Figure 5.**
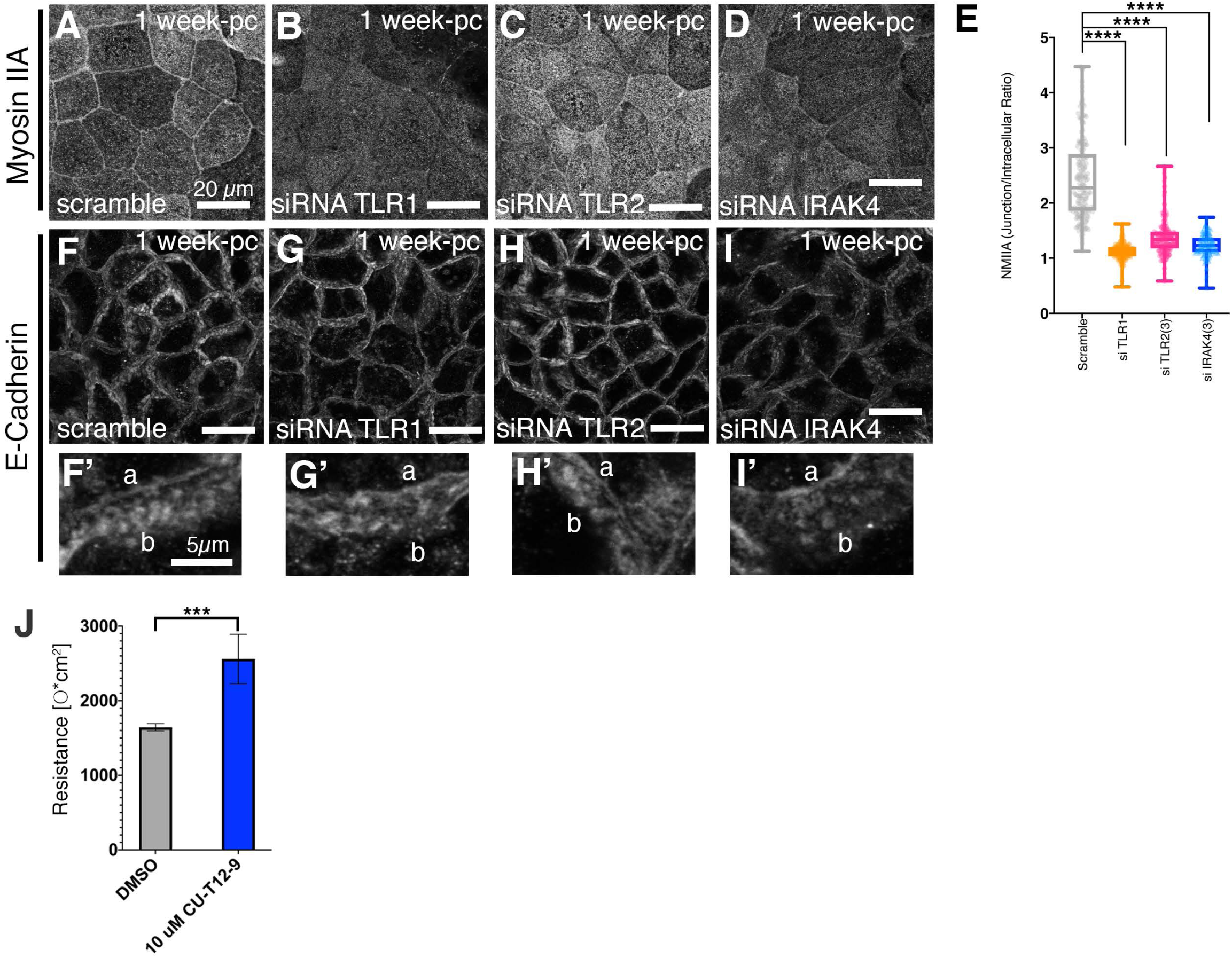
Reduction of TLR levels leads to loss of epithelial tightness and changes at junctions. In siRNA-treated 1-week post-confluence Caco-2 monolayers with reduced TLR1 or TLR2 levels, localisation of junctional components to cell borders is affected. Z-projections of confocal sections covering the apical-lateral junctional area are shown. **A-E** Knock-down of TLR1, TLR2 and IRAK4 leads to MNIIA reduction at junctions. **E** Quantification of effects: n= 182 junctions were analysed for the Scramble control and n= 142 for siTLR1 and n=187 for siTLR2 and n= 151 for siIRAK4. Statistical significance was determined by one way ANOVA with Dunnett’s multiple comparison test as **** = p<0.00001. **F-I’** Knock-down of TLR1, TLR2 and IRAK4 leads to changes to E-Cadherin distribution at lateral junctions, similar to IRAK4-inhibitor treatment. **F’-I’** show higher magnifications of individual tilted lateral sides of Caco-2 cells, ‘a’ indicates the apical and ‘b’ the basal end of the lateral membranes shown. **J** Treatment of Caco-2 cells with 10µM of the TLR1/2 agonist CU-T12-9 leads to an increase in TEER after 5 days in culture (upon seeding at confluency). Error bars indicate SD of n= 3 (DMSO, CU-T12-9) separate transwells. Statistical significance was determined by unpaired Student’s t-test as *** = p<0.0001.

## Acknowledgments

The authors would like to thank A.Nagafuchi for the α18-α-Catenin antibody. We thank members of the lab for critical reading of the manuscript.

The work is supported by the Medical Research Council (file reference number U105178780) and the LMB-AstraZeneca Blue-Sky Fund (BSF21).

## Author contribution

Conceptualisation, K.R. and J.P..; Methodology, K.R., J.P, A.B., K.B.S..; Investigation, K.R., J.P.; Writing-Original Draft, K.R., J.P.; Funding Acquisition, K.R., A.B., K.B.S.

